# GEGA (Gallus Enriched Gene Annotation): an online tool providing genomics and functional information across 47 tissues for a chicken gene-enriched atlas gathering Ensembl & Refseq genome annotations

**DOI:** 10.1101/2024.03.13.584813

**Authors:** Fabien Degalez, Philippe Bardou, Sandrine Lagarrigue

**Author notes:** Joint Authors (Fabien Degalez, Philippe Bardou).

## Abstract

GEGA is a user-friendly tool to navigate through different genomics and functional information related to an enriched gene atlas in chicken that unifies the gene catalogues from the two reference databases, NCBI-RefSeq & EMBL-Ensembl/GENCODE, and four other additional rich resources as FAANG and NONCODE. Using the latest GRCg7b genome assembly, GEGA offers a total of 78,323 genes, including 24,102 protein-coding genes (PCGs) and 44,428 long non-coding RNAs (lncRNAs), greatly enhancing the number of genes provided by each resource separately. But GEGA is more than just a gene database. It offers a range of features that allow to go deeper into the functional aspects of these genes, *e*.*g*., by exploring their expression and co-expression profiles across 47 tissues from 36 datasets and 1400 samples, by discovering tissue-specific variations and their expression as a function of sex or age, by extracting their orthologous genes or their configuration related to the genomics closest gene. For the communities interested in one specific gene, a list of genes or a QTL region in chicken, GEGA’s user-friendly interface enables efficient gene analysis, easy downloading of results and a multitude of graphical representations, from genomic information to detailed visualization of expression levels.

**GRAPHICAL ABSTRACT:** 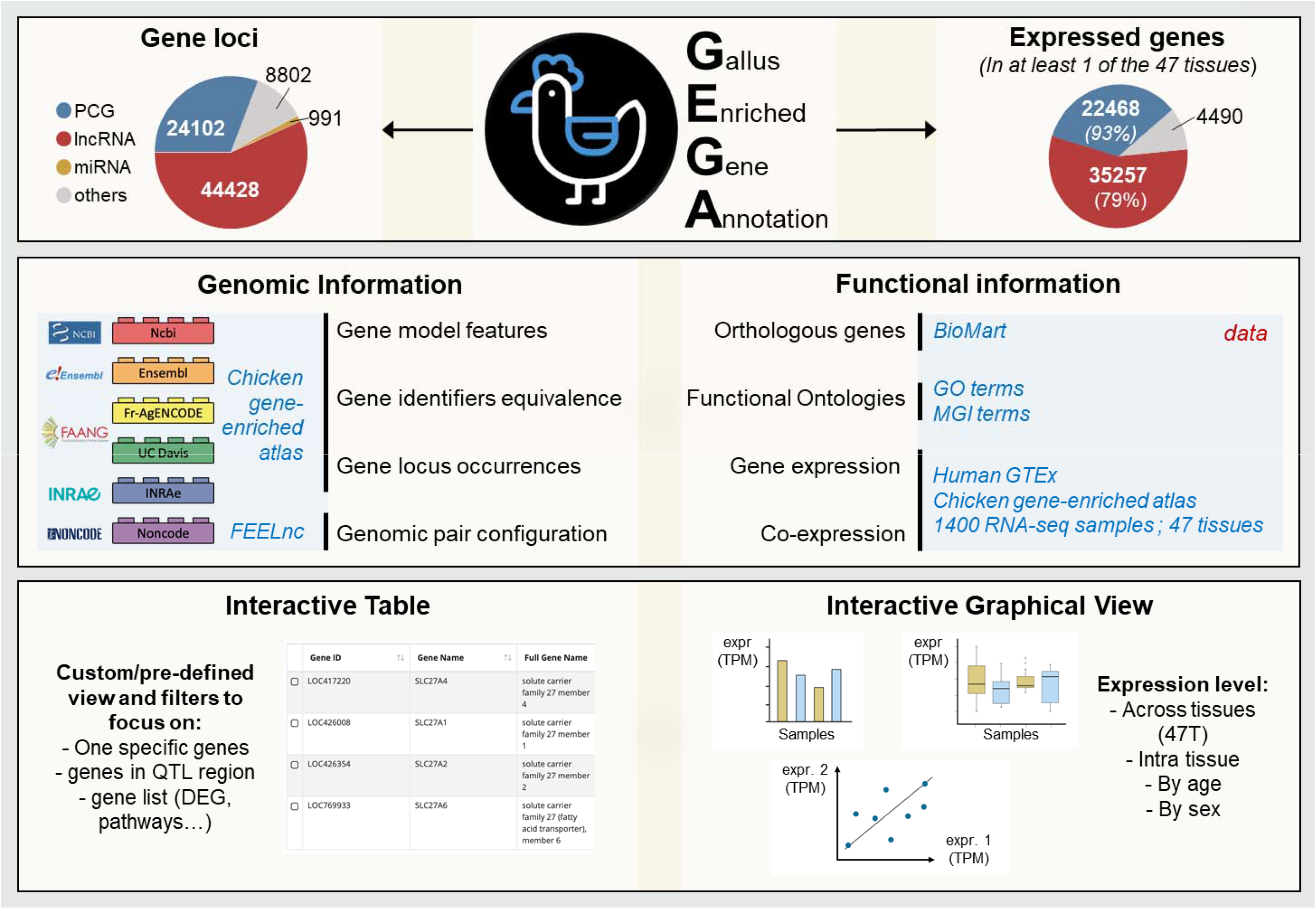

## INTRODUCTION

The chicken (Gallus gallus) genome is a valuable model in both fundamental and applied research (1). This key model species is often used in order to investigate vertebrate development, evolution and more generally diseases. Moreover, the chicken is one of the livestock species which supplies the most protein-rich food worldwide through both meat and egg production. In recent years, significant progress has been made in annotating this genome with, *e.g*., the introduction of long non coding RNA (lncRNA) gene models in the two reference gene model databases, NCBI-RefSeq and EMBL-EBI Ensembl/GENCODE. These two databases, which are widely recognized as reputable resources, provide complementary information concerning gene models due to different RNAseq resources and bioinformatics pipelines used for the gene modelling (2). As previously reported (3), the overlap of protein coding gene (PCG) models between both reference databases is around 90% with, generally, different transcript models supporting the common PCG loci. Concerning lncRNA gene, which are regulatory elements of gene expression characterized by condition-specific expressions (tissue and temporal), making them harder to identify, the overlap between their loci reaches around 25% between both annotations. By combining the two reference annotations and providing gene identifier correspondence for the common gene loci, the scientific community can exploit the strengths of both databases, providing access to a greater number of gene models and improving the accuracy of their analyses while easily using the tools provided by each database. However, even if some initiatives such as the MANE project for human (4), aims to determine common gene models between these annotations, to date and to our knowledge, an annotation resulting from the union of these reference databases is not currently accessible in an easy and systematic way. In this context, we recently released an updated gene atlas of the chicken genome based on the last GRCg7b assembly, integrating reference annotations from both “NCBI-RefSeq” and “EMBL-EBI Ensembl/GENCODE” databases and other gene models from additional resources as multi-tissue projects initiated by the FAANG consortium (5–7) or lncRNA dedicated databases as NONCODE (8) as presented in Degalez et al., 2024 (3). Briefly, this gene atlas consists of 78,323 gene models including 24,102 PCGs and 44,428 lncRNAs with a total of 63,513 (81%) genes considered as expressed (≥ 0.1 TPM) including 22,468 (93%) PCGs and 35,257 (79%) lncRNAs and 43,252 (55%) considered as highly expressed (≥1 TPM) including 19,819 (82%) PCGs and 20,252 (46%) lncRNAs. Indeed, all these genes were functionally annotated through their expressions across 47 tissues using 1400 RNA-seq samples from 36 dataset chosen to represent the diversity of chicken physiological systems. This gene atlas eases switching between “EMBL-EBI Ensembl/GENCODE” and “NCBI-RefSeq” gene identifiers, as well as between assembly versions, since it provides gene identifier equivalents for the previous galgal5 and GRCg6b assemblies of the chicken. This gene loci resource is also complemented by various functional annotations such as orthologous genes in human and mouse, gene ontologies (functional Gene Ontology terms or phenotype terms) or by configuration with the nearest genes

To easily access and explore all this information (gene models, expression information and the associated functional annotations), we have developed an online tool called GEGA (Gallus Enriched Gene annotation). GEGA is addressed to different communities, whether those interested in genomic regions associated with phenotypes of interest (QTL) or those working directly on gene expression and interested by one specific gene or a list of genes. In this paper, the GEGA tool and the different usages are described.

## MATERIAL AND METHODS

### Webserver Infrastructure

The system is built on an Apache web server, version 2.4.37, configured with OpenSSL version 1.1.1k to ensure communication security. The database server is powered by MariaDB, version 10.3.35-MariaDB for data storage and retrieval. This software configuration was deployed on a Rocky Linux distribution.

### Back-End Development

The back-end development was carried out using Node.js with the Express.js framework. Data is stored in a MySQL database, with a specific instance of MariaDB version 10.3.35 chosen for its compatibility and enhanced performance. These technologies enabled efficient data manipulation and optimal handling of HTTP requests.

### Front-End Development

Bootstrap, version 4.6.0 was selected as the primary CSS framework for the development of the user interface, ensuring a responsive and aesthetically pleasing design for an enhanced user experience across different devices. For dynamic data manipulation and the creation of interactive tables, jQuery, version 3.5.1, and DataTables, version 1.10.16 were used.

### Data Visualization

Regarding data visualization, Highcharts, version 10.3.3, a powerful and flexible JavaScript library, allowing the creation of interactive and customizable charts was used. This integration provided a clear and intuitive visual representation of the results generated by data analysis.

*Origin and processing of data used in GEGA are detailed in a more exhaustive way in the associated paper, Degalez et al*., *2024 (3). Only a summary is provided here*.

### Reference assembly

The genome annotation is established on the bGalGal1.mat.broiler.GRCg7b (GCF_016699485.2) assembly of the chicken (Gallus Gallus) genome (9).

### Data origins – Individual database

The enriched genome annotation included in GEGA integrates, as detailed in Degalez et al. 2024 (3), gene models from six sources: the “NCBI-RefSeq” (v106) (10) and “EMBL-EBI Ensembl/GENCODE” (v107) (11) reference databases according to GRCg7b assembly and based on around 300 (12) and 60 RNA-seq (13) samples respectively; the “FR-AgENCODE” (14) and “FarmENCODE” (15) FAANG pilot annotation projects (5–7) produced under GRCg6a assembly; the gene annotation from Jehl et al. 2020 (16) according to the galgal5 assembly and integrating gene models from different public databases complete by 364 new RNA-seq samples; and, finally, the NONCODE annotation (8) including non-coding gene models under the galgal4 assembly. Note that, if the FarmENCODE project used Oxford Nanopore long-read sequencing, the five other projects mainly used short-read RNA-seq data. Gene models from assemblies older than GRCg7b were remapped using the NCBI genome remapping service with default parameters (17). Quality of gene models was evaluated by the overlap degree of transcripts with FANTOM5 CAGE peaks (18) remapped from galgal5 to GRCg7b. Considering also the database popularity, gene models were added sequentially resulting in this order of aggregation: 1) RefSeq; 2) Ensembl; 3) FrAg; 4) Davis; 5) Inrae; 6) Noncode. Gene models were added from each database to the growing atlas, if their associated transcripts did not overlap genes already present in the growing atlas. Two genes were considered as overlapping if one of their transcripts shared at least one exonic base pair on the same strand using BEDtools (19). To limit overlapping similar patterns with different biotypes, models were aggregated by biotype class.

### Data origins – Expression

A set of 36 datasets publicly available including a total of 1400 samples were chosen to represent a variety of 47 tissues. The list of the 47 tissues, their abbreviations and color codes used are available on GEGA. For each tissue in each project, a median of TPM normalized expressions across samples was calculated. For tissues present in several projects, the median was calculated using the TPM medians previously calculated in each project. A gene was considered as expressed if *i)* its median expression was ≥ 0.1 TPM in at least one tissue, and *ii)* at least 50% of samples of a tissue for a given project have a reads number ≥ 6 as well as the normalized TPM and TMM expression ≥ 0.1.

### Tissue-specificity analysis

Because the tissue specificity is very sensitive to the chosen indicator, different indicators of tissue specificity are available in GEGA:

*i)* the tau metrics, assessed using raw (τ) or log10 median tissue expression (τ_log10_) in TPM (20), and defined as follows:

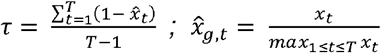

*with x*_t_ *the raw or log10 median expression of the gene of interest in the tissue* t *and among the* T *tissues*.

A gene was considered as tissue specific for a τ or τ_log10_ ≥ 0.90 ; the columns “expr_isTS” or “expr_isTSlog10” are then equal to 1. Therefore, even if the gene is considered as tissue specific, its expression profile can vary considerably and show different breaks. A break was defined as a difference in expression by a factor of 2, *i.e*., FC ≥ 2 when expressions were ranked in descending order.

The number of observed breaks, associated FCs, and tissues in each subgroup are indicated in the columns “expr_nbBreaks”, “expr_fcBreaks” and “expr_tissueBreaks” respectively. *ii) the* FC between the first and second most expressed tissues or *iii)* the FC between the two first groups separated by a break, that was calculated in three different ways, considering: *a)* the least expressed tissue in the first group and the most expressed tissue in the second one (“expr_fcFirstBreak_lowToTop”); *b)* the most expressed tissue in both groups (“expr_fcFirstBreak_topToTop”); c) the median expression for both groups (“expr_fcFirstBreak_medianGroup”).

### Classification according to the closest feature

PCG, lncRNA, miRNA and small RNA (smlRNA including snoRNA, snRNA, sRNA, tRNA, rRNA, scaRNA) transcripts and genes were classified through their associated transcripts relatively to their closest PCG and lncRNA transcript using FEELnc v.0.2.1 (default parameters, 100kb between the TSS of the transcripts of the gene of interest and the transcripts of its closest lncRNA or PCG gene model) (21). Briefly, gene pairs are split into two categories and three sub-categories. Firstly, the gene of interest in the pair is considered to be “genic” if it overlaps the partner gene, and “intergenic” otherwise. Secondly, the gene of interest is classified according to its configuration with its partner: “same strand / sense” if it is transcribed in the same orientation; “divergent / antisense” if it is transcribed in head-to-head orientation and; “convergent / antisense” if it is oriented in tail to tail. For each type of pairs (*e.g*., lncRNA:PCG noted ‘LncPcg’ in which the lncRNA is the gene of interest), different information are provided such as the class name of the configuration according to FEELnc (*e.g*., lncgSSinNest, which means that the lncRNA gene is nested in the intron of the PCG), the distance between the two genes, or also the gene id and the gene name of the closest gene (here the PCG) of the gene of interest. For more details, see the columns description available on GEGA (hyperlink associated to “show/hide columns”).

For each lncRNA:PCG, lncRNA:lncRNA and PCG:PCG pairs for which the tissue expression were available, the Kendall correlation (τ) between the expression values across tissues was computed, providing a co-expression indicator between the two genes of the pair.

### Orthology and GO terms

Gene orthology between the chicken, the mouse and the human as well as GO terms and phenotype description were extracted using BioMart (22) from Ensembl (V107). GO terms come from the “Gene ontology” database (23, 24), one of the most complete and widely used “functional” databases. Phenotype description refers to the OMIA database (25). The Ensembl reference database chosen for extracting this information allows us to maintain it easily via the “biomart” API tool at each update.

### GTEx data analysis

A list of 23 equivalent tissues between the human and the chicken has been established and the abbreviations and color codes used are available on GEGA. The median gene level TPM for these 23 tissues from RNA-seq data of human GTEx Analysis V8 was used (26).

### Summary of the available information

For each gene, a total of 170 information is available and described in the columns description available on GEGA (hyperlink associated to “show/hide columns”).

## RESULTS

### General description of the GEGA tool

GEGA is an online tool enabling the exploration of an atlas of 78,323 gene models including both genomics and functional annotations according to the chicken GRCg7b genome assembly. Three related main features are available (Figure 1): *(i)* an interactive table, in the upper part of the “Explore” menu, with possibilities to view different gene features as presented briefly in the “Mat. and Met.”; *(ii)* an interactive viewer, in the lower part of the “Explore” menu, providing plots to explore the different patterns of expression and co-expression for gene models according to different conditions (inter tissues / intra tissue; sex / age) but also to explore gene models information previously selected; (iii) an interactive viewer in the “Browse” menu to visualize gene models according to the six resources used for generating the gene enriched atlas. This tool is publicly available on https://gega.sigenae.org/ and free to use. It does not depend on cookie files or any credentials. The web server was tested on Chrome V.116.0, Firefox V.116.0 and Safari V16.5.2. browsers. A dedicated contact tab is available to help improve the tool, particularly if ideas for improvement, conception problems or bugs are identified.

**Figure 1.**
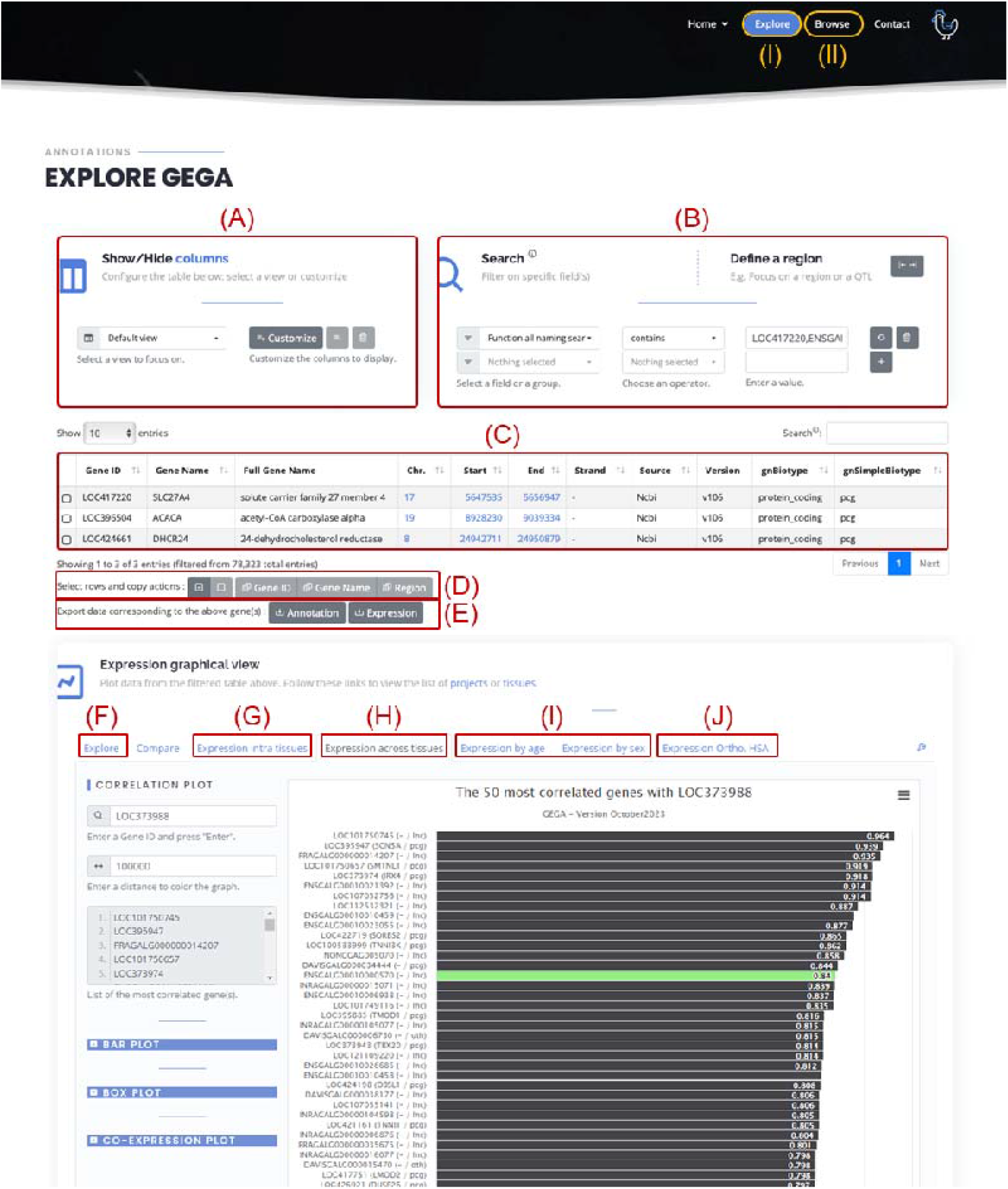
Interface overview. (I) GEGA main page to select genes of interest and observe their expression, including: (A) View selection panel; (B) Filter selection panel; (C) Table of selected gene according to the chosen view; (D) Copy panel; (E) Exportation of annotation and/or expression; (F) Visual exploration of the selected genes; Graphical tab for (G) intra-tissue, (H) inter-tissues, (I) age/sex and (J) human ortholog expressions. (II) Access to the browser to view the enriched atlas and the six annotations used to build it.

### Interactive Table – Data viewing

A total of 170 information columns are available (see “show/hide column link” in GEGA interface), but the user may not be interested in all of it. Therefore, only data concerning the gene model *stricto sensu* are initially displayed by default. The user can display additional data through the dedicated “Views” panel (Figure 1A). Pre-filtered views were implemented to guide potential new users or facilitate access to specific information, following the main categories of information available in GEGA. The first three columns, which are always identical, include the gene identifier from the source and the short and long gene names. The preconfigured views are as follows: *(i)* “Default”, displaying the origin, positional information, and biotype; *(ii)* “All naming”, showing available identifiers for a gene model, particularly Ensembl and NCBI, and for previous assemblies including galgal5 and, GRCg6a; *(iii)* “All functional”, listing gene ontology (GO) and phenotype terms for each gene and their associated orthologs; *(iv)* “Expression & Tissue-specificity”, indicating expression values (in TPM) in each of 47 tissues, top tissues, and tissue specificity; *(v)* “Orthology”, showing orthologous relationships and equivalents with human and mouse for each protein coding gene; *(vi)* “Gene structure”, detailing transcript, exon, and intron structure; *(vii)* “FEELnc” indicating the nearest gene model (PCG/lncRNA/miRNA) and configuration at the gene and transcript level, along with co-expression data at the gene level; *(viii)* “Repetability”, providing reproducibility information on loci representation across resources. For flexibility and specific usage, columns can be manually selected with the “Customize” tool using names or the tree structure with checkboxes.

### Interactive Table – Data filtering

Filters can be easily applied to extract a specific set of genes using the “Search” panel (Figure 1B). All columns available in the “Views” panel can be filtered and combined. Some filters can be applied simultaneously using predefined “custom function” available in the same selection menu. To date, the “Function all naming search” and “Function all functional annot. search” allow to filter across all the naming (names and gene identifiers from both Refseq and Ensembl) or GO related columns respectively. Logical operators can be applied according to the data type. The “equals/not equals” operator can be used for both numerical and categorical data, while “greater/lower” is specific to numerical data and “contained/not contained” to categorical data. For a given filter, the comma (“,”) delimiter can be used to indicate several possibilities (OR operator), *e.g*., application of the filter “gnSimpleBiotype – contains – lnc,pcg” will results of a selection of both lncRNA and PCG. In the same way, “Function all naming search – contains – LOC417220,ENSGALG00010025549,ACACA” will result in a selection of the three genes mentioned, even if the type of nomination used varies. Note that, each line corresponds to a criterion and a set of criteria (AND operator) is considered as an intersection of condition. Even if each tissue expression is not available individually through the interactive table, filter on tissue gene expression is available through the “[tissueName] median” filter. Note that filters defined in the “search” panel apply even if the column is not selected in the “Show/Hide columns”.

To answer to specific usage (see user cases), a “Define a region” module to create a region surrounding a gene (using identifier or name) or position (defined by chromosome and position) has been designed and can be used, *e.g*., for QTL analysis. Note that offsets can be applied equally or differently on each side.

### Interactive Table – Output data

After filtering, genes of interest (*i.e*., lines of the table – Figure 1C) can be easily selected and either gene names, gene identifiers or the genomic region can be copied to the clipboard (Figure 1D). The selected list of genes or the genomic region can be then paste elsewhere for further analysis, in particular for interactive visualization. Two types of filtered tables are available for download (Figure 1E) as a .*csv* file depending on user preference: *(i)* “Annotation”, which includes all available information or only what is displayed through the custom view as chosen by the user ; (*ii*) “Expression” which provides tissue expression levels for the filtered genes in the 47 tissues.

### Interactive visualization – Explore genes according to the filtered interactive table

An interactive visualizer provides a graphical overview of gene sets selected by specific filters (Figure 1F). Consequently, pie charts and boxplots can be generated for categorical and numerical variables, respectively. Considering two numerical variables, a scatter plot showing potential relationships between them can be plotted, *e.g*., lncRNA and PCG co-expression *vs*. separating distance.

### Interactive visualization – Global usage to explore the gene expression

Expression patterns can be analyzed either across 47 tissues or within a specific tissue in which variation between individuals and projects can be observed. It is also possible to observe gene expression according to factor such as the sex (projects “FrAgENCODE”:n=2 (14) and “RIRinrae”: n=4). or ages (projects “Kaessmann”:n=4 (27) and “RIRinrae”:n=4) in which multiple tissues have been studied. Although each feature has its own specificity, boxplots and co-expression plots (scatter plots) are always available and work similarly. For boxplots, a gene or list of genes (separated by spaces or commas) can be provided. Expression for each gene will be displayed in the input order. To enable comparison, expression can be unified by displaying all values on a scale of 0 to the maximum observed across all genes. Sample identities (individuals) can also be displayed. Tissues are classified by abbreviation by default, but can be ordered by expression proximity as analyzed in Degalez et al., 2024 (3). Tissues sharing color ranges generally share a physiological system. For co-expression plots, a reference gene identifier is first provided, followed by a target gene(s). Each plot of the reference (X-axis) versus target (Y-axis) gene is displayed, initially in input order. However, plots can be ordered by correlation (1 to -1) or filtered by a correlation threshold. Because co-expression can follow different laws, linear or logarithmic plots are available. Displaying the regression is also an option.

### Interactive visualization – Specific usage: “Expression intra tissues” (Figure 1G)

First, after selecting the tissue of interest, expression profiles across 47 tissues can be displayed for the 40 most highly expressed genes to identify tissue-specific or high-expressed ubiquitous genes. Second, boxplots of expression for each project related to the chosen tissue or co-expression plots between gene pairs of interest across all samples of the all projects related to the chosen tissue can be displayed.

### Interactive visualization – Specific usage: “Expression across tissues” (Figure 1H)

First, to overview gene expression correlation across 47 tissues, a correlation plot is available for one gene of interest. Once the gene identifier has been provided, the 50 most correlated genes are displayed in descending order of correlation. The gene list is accessible and can be copied/pasted for further investigation. Note that genes on the same chromosome as the gene of interest appear in blue, and in green if their distance is less than the specified threshold. This can help identify clustered co-expressed genes, regardless or not of chromosomic location. As an example, TBX5 (LOC373988; chr15:12,317,331) shows a high co-expression across tissues (ρ=0.84) with a lncRNA (ENSGALG00010000570; chr15:12,314,252) which is in 15th position and colored in green because both genes are separated by less than 100kb (default value).

For examining expression across tissues, after providing a gene or a gene list, either a boxplot (as described before) or barplot of expression across the 47 tissues can be generated. Boxplots consider all samples from all projects together, plotting all available data. Barplots first calculate the median expression for each project, then the median of those medians, thus lacking sample resolution. Barplots have the same options as boxplots.

### Interactive visualization – Specific usage: “Expression by age/sex” (Figure 1I)

First, a project of interest must be selected among the few projects related to multiple ages (projects “Kaessmann” and “RIRinrae”) or sexes (projects “FrAgENCODE” and “RIRinrae”). This ensures reliable and consistent sub-group comparisons across studied factors. Then, the tissue of interest has to be selected among the tissues available for the selected project. Finally, barplots of gene expression across samples can be generated according to the factor levels (ages or sexes) on linear or logarithmic scales, with unified or raw maximum values. Boxplots can be generated for one specific tissue as barplots, but also for different tissues, up to four tissues. Boxplots can be then ordered by condition or tissue depending on what the user wants to highlight. Finally, co-expression plots using all available samples for a given tissue of a selected project can be displayed, following the previously described process.

### Interactive visualization – Specific usage: Expression Ortho. HSA (Figure 1J)

For chicken gene with a human orthologue, expressions in both species can be observed across 22 equivalent tissues. Indeed, after inputting the chicken gene identifiers, the associated one-to-one human orthologs are retrieved and both expressions can be displayed either *i)* through a stacked barplot for a side-by-side comparison with the possibility of grouping tissues according to their functional proximity, as defined previously, or *ii)* through a coexpression plot, to observe overall expression patterns. For a given gene, the raw expressions values for both species can be download via a .*csv* or .*xls* file.

### Interactive visualization – Output data

Regardless of plot type, the associated figure can be downloaded in PNG, JPEG, SVG, or PDF format through the dedicated menu (top-right of plot). The data used for plotting can be downloaded as a .*csv* formatted to enable re-plotting with specific tools (*e.g*., R-base or ggplot). Following this principle, production of multi-plots is currently unavailable; each plot and associated data must be downloaded independently.

### Interactive Browser

The “Browse” menu displays a genomic region or gene of interest along with the surrounding genes at the chosen resolution. It is possible to observe the gene models present in the enriched atlas, but also in each of the six resources used to generate it, indicated by a specific color. The gene models can be observed with their specific orientations as a classical genome browser and therefore visualize the FEELnc gene pair configurations indicated in the interactive table. For example, we can observe TBX5 (LOC373988; chr15:12,317,331) in divergent orientation with its lncRNA (ENSGALG00010000570; chr15:12,314,252).

### GEGA usages through typical user cases

#### Focus on tissue-specific (TS) analysis (Figure 2)

For genes of interest, depending on the dataset used and the metrics applied, the tissue-specificity (TS) can provide additional information. In GEGA different metrics are available. *i)* The tau metrics, τ: “expr_tau” with raw expressions or τ_log10_: “expr_tauLog10” with expressions in log10, and “expr_isTS” and “expr_isTS_log10” indicating a τ and τ_log10_ ≥ 0.90 respectively. These tau metrics provide a unique score relatively to the expression pattern of the 47 tissues. *ii)* The fold changes (FC) between the two most expressed tissues (“expr_fcTop1Top2”). Note that a τ ≥ 0.90 corresponds to an average and median FC of 71 and 2.5 for tissue with expression ≥ 1 TPM in at least one tissue and 5 and 2.19 for tissue with expression ∈ [0.1,1 [TPM. However, as already mentioned, the τ and FC metrics correspond to a unique tissue, but a gene can be TS for several tissues which are functionally close. Therefore, the concept of “break” was introduced, allowing distinguishable tissue groups to be separated according to the expression pattern. The list of tissues and the total number included in the first group can be observed using the “expr_tissueFirstBreak” and “ expr_tissueNbFirstBreak “ filters respectively. iii) Three metrics have been defined to quantify the expression fold change between both groups delimited by the first break (“expr_fcFirstBreak_lowToTop”, “expr_fcFirstBreak_topToTop”, “expr_fcFirstBreak_medianGroup”; see Mat. and Met. for details). As an example, results of each metrics on both expressed PCGs and lncRNAs in the liver are available in Figure . As expected, lncRNA genes are more TS than PCGs as previously reported in chicken (3, 16) and other species (28). As illustrated for PCG in the liver, note that each TS metric possesses its own characteristics, with a variable number of TS genes being shared by several methods. As an example, the 90 TS genes detected by the “FC” metric are included in the “tau_livr” metric (n = 205), which itself is 94% included in the “FC1st” metric (n = 264). For this latter, it corresponds to 3 TS numbers according to the manner in which the FC is calculated between groups (n = 125, n = 163, n = 141). The intersection results between each TS metric can be observed via Venn diagrams produced using the “Compare” tab available in the graphical view of GEGA or in Supplementary Figure 1 for this example.

**Figure 2.**
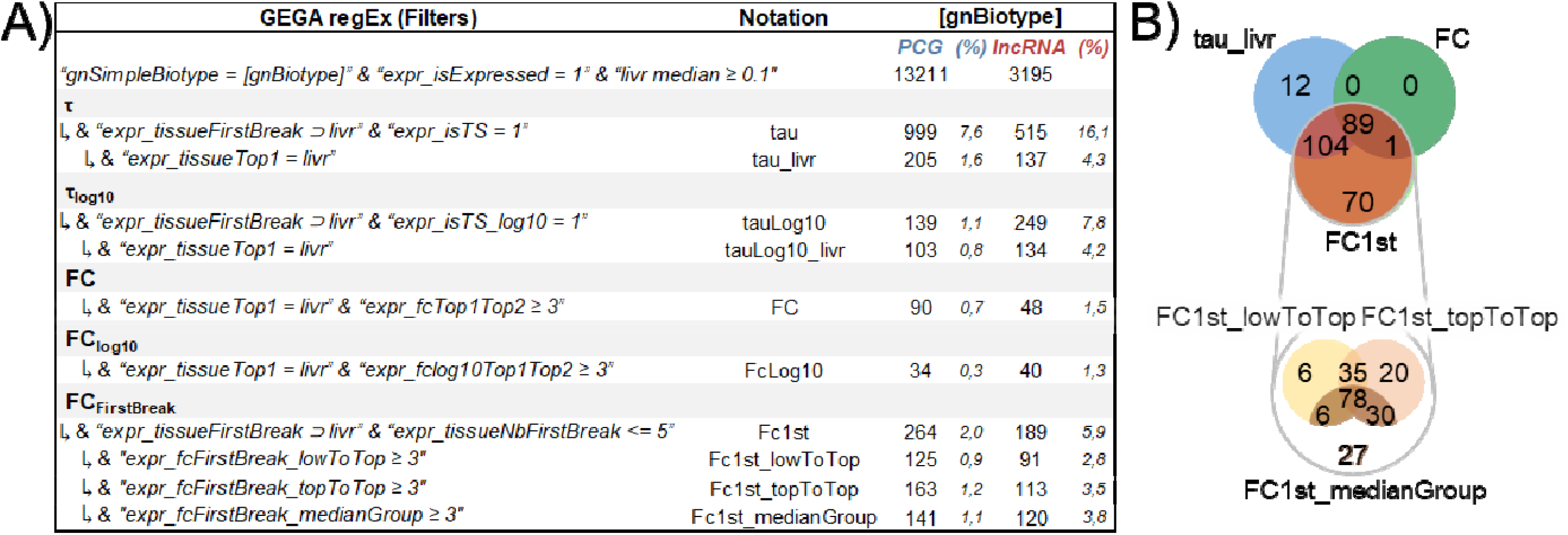
Tissue-specific metrics. (A) Number of PCGs (blue) and lncRNAs (red) considered as tissue-specific in the liver according to the various metrics available in GEGA. (B) Intersection of the TS metrics for PCGs in the liver.

#### Focus on one specific gene (e.g., TBX5; Figure 3)

TBX5, a T-box transcription factor (29), plays a critical role in vertebrate cardiac morphogenesis. Overexpression of TBX5 in embryonic chick hearts has shown that TBX5 inhibits myocardial growth cardiogenesis and thereby participate in modulating vertebrate cardiac growth and development (30). Using GEGA, this gene can then be deeply explored, *e.g*., following this process: 1) Analysis of expression across the 47 tissues and 2) through functional terms (GO terms / phenotype terms) to underline those which are consistent with the TBX5 function. 3) Observation of genomics configuration between the two neighboring genes. This feature underlines the presence of an antisense lncRNA. 4) Gene orthology examination for human and mouse. The TBX5 gene has an orthologous PCG in mouse and human. This initial observation served manual verification of the conservation in both species of the antisense lncRNA. 5) Analyses of expression and co-expression across the 47 tissues. TBX5 and its antisense lncRNA revealed that the lncRNA is highly expressed in heart as TBX5. Moreover, both genes are highly co-expressed in heart samples. 6) Analyses of expression related to ages. The gene pair is more expressed in embryo than adult stages.

**Figure 3.**
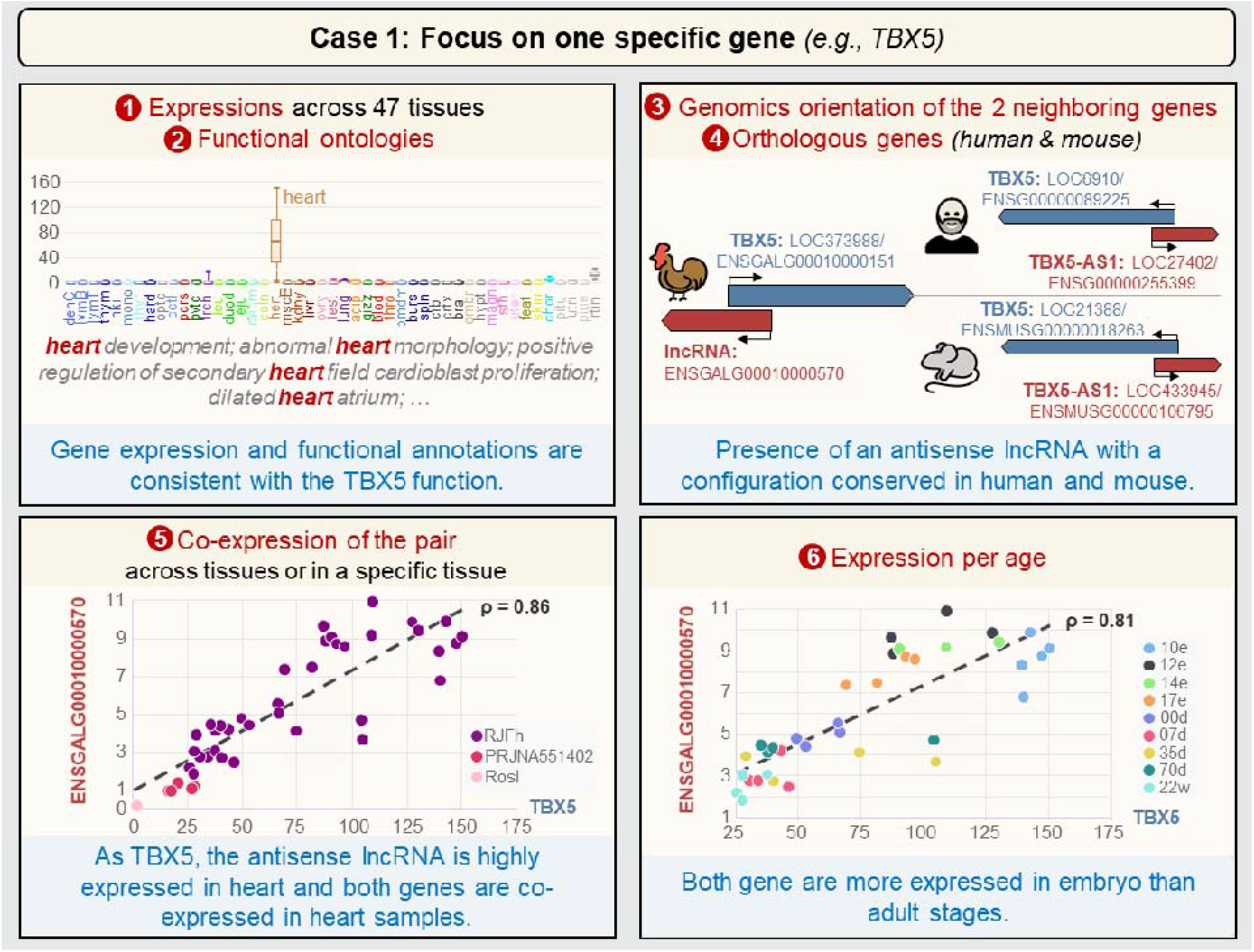
User case 1 – GEGA for exploring a specific gene, its genomic environment and its potential functions (e.g., TBX5). Correspondence between full tissue names and *the* 4 letters abbreviations is available on GEGA (hyperlink associated to “tissues” in the “Expression Graphical View”). *Additi*onal project-specific information is available on GEGA via the hyperlink associated with “projects” in the “Expression Graphical View”. e: embryonic day; d: day; w: week.

This data exploration by GEGA suggests that the antisense lncRNA (known as TBX-AS1 in human) could be a regulator of TBX5 or at least could share a common function. This result is consistent with the fact the cardiac transcription factor gene (TBX5) is accompanied by a bidirectional long non-coding RNA as reported in 2018 by Hori et al. (31).

#### Focus on a QTL region (e.g., chicken epilepsy; Figure 4)

A QTL for epilepsy trait in chicken was mapped in 2011 around the markers 100A3M13 and SEQ1009, that mapped at that time to linkage group E26C13 which has been identified thereafter as the micro-chromosome GGA25 (32). Through fine mapping and other molecular approaches, the authors identified the SV2A gene as related to the epilepsy trait. GEGA could have facilitated the identification of this gene as follows: 1) Definition of a region of +/- 250kb around 100A3M13 (*i.e*., chr25 from 641442 to 641643) and display of all the genes, with the possibility of filtering the biotype of the genes (protein-coding genes (PCG), long non-coding genes (lncRNA), …). In the current atlas, 39 protein-coding genes are observed in this region. 2) Analyses of the functional terms of these 39 genes. Of these, 8 genes respond to GO terms or phenotypes associated with the trait of interest, *i.e*., “brain,neuron,synapse,epilepsy”. 3) Analyses of the expression across the 47 tissues. Only SV2A is specifically expressed in the cerebral system, which is consistent with the trait of interest. 4) Observation of the orthologous gene expression in human shows that the expression pattern is conserved between both species.

**Figure 4.**
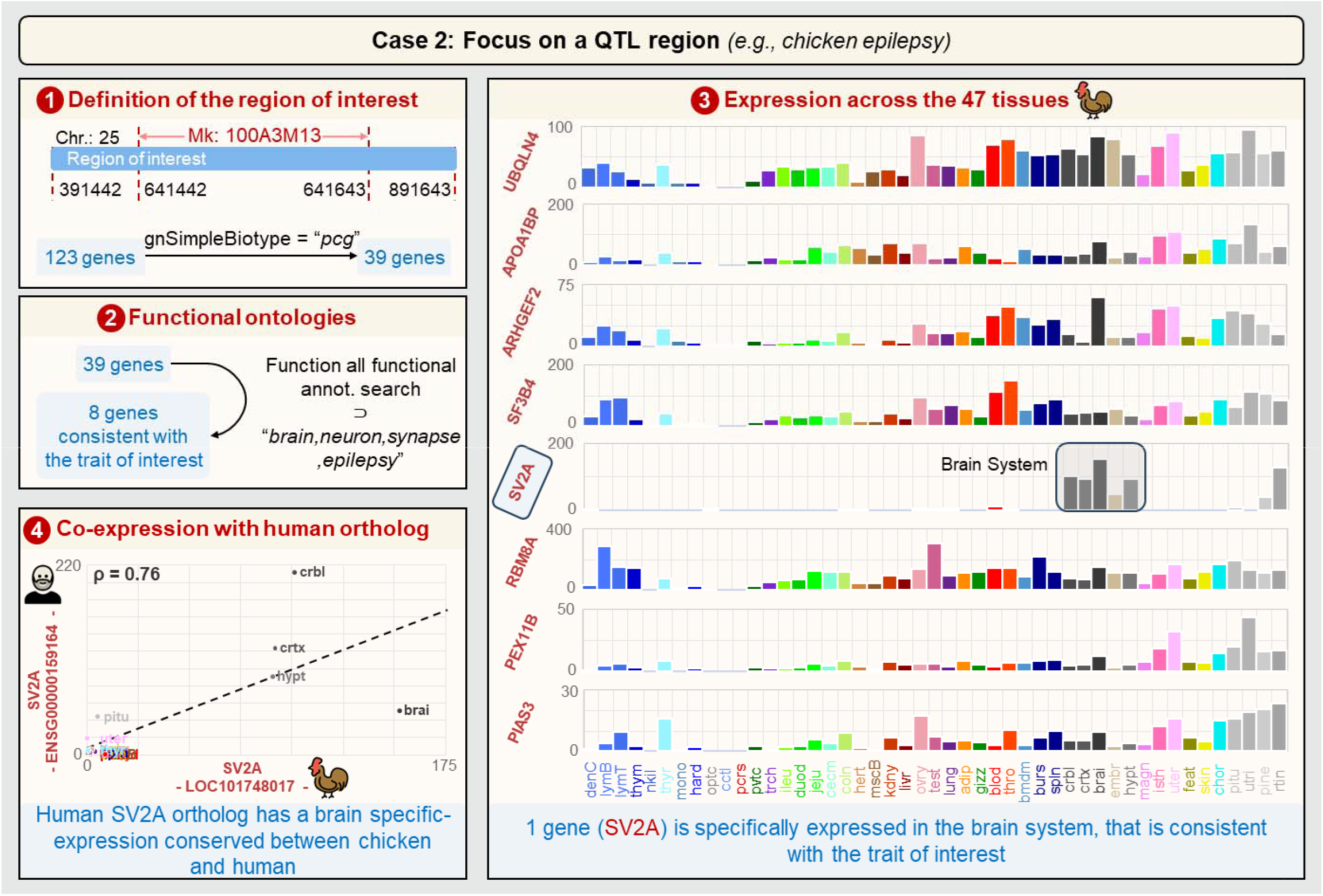
User case 2 *–* GEGA to analyze a QTL region (e.g, epilepsy region around the marker 100A3M13). Correspondence between full tissue names and the 4 letters abbreviations is available on GEGA (hyperlink associated to “tissues” in the *“*Expression Graphical View”). Chr.: Chromosome.

#### Focus on a gene list (e.g., fatty acid synthesis and transport genes; Figure 5)

The main role of liver in the chicken adaptive response to a switch in dietary energy source through the transcriptional regulation of lipogenesis was previously reported. Indeed, 298 down-regulated genes in “High Fat / High Fiber diet (HF)” compared to a standard diet containing “Low Fat / High Starch” (LF) was observed with an enrichment of genes related to fatty acid synthesis and transport (33). The expression in liver of TADA2A (Transcriptional Adaptor 2A) was highly correlated to gene expression related to key enzymes of fatty acids (FA) anabolism as ACACA, FASN, SCD, DLAT, MTTP and ELOVL6. Thanks to these results, TADA2A has been identified as a potential new player in the regulation of lipogenesis. Analysis of the co-expression of TADA2A with these six genes of interest can be further investigated using GEGA. Using the 11 projects related to liver with 265 samples, the hepatic co-expression of TADA2A (LOC417656/ENSGALG00010029362) with ACACA, FASN, SCD, MTTP, DLAT and ELOVL6 (LOC396504/ENSGALG00010029360, LOC396061/ENSGALG00010029360, LOC395706/ENSGALG00010021083, LOC422709/ENSGALG00010005268, LOC419796/ENSGALG00010025156, and LOC428772/ENSGALG00010003790 respectively) is confirmed (0.65 ≤ ρ ≤ 0.87), with the highest correlations for ACACA and FASN (ρ ≥ 0.83). The co-expression in brain was then explored since fatty acids are essential for brain functions (34) and a co-expression between TADA2A and ACACA in mouse brain was already reported (33). In the hypothalamus (2 projects), TADA2A is highly co-expressed with ACACA, FASN, DLAT, ELOVL6 but not with SCD & MTTP (ρ ≤ 0.15). Co-expression in the brain as a function of age (through the “Kaessmann” project) shows that TADA2A is highly co-expressed with ACACA and FASN across embryo and adult ages; TADA2A is positively co-expressed across adult ages and negatively co-expressed across embryo ages with DLAT and ELOVL6; it is not co-expressed with SCD and MTTP. In summary, GEGA shows that, TADA2A appears highly co-expressed with ACACA & FASN genes coding the two first key enzymes of lipogenesis, whatever the projects, the ages and the analyzed tissues (liver or brain/hypothalamus) whereas the co-expression pattern is multi-form across these different conditions/tissues with DLAT, ELOVL6, SCD.

**Figure 5.**
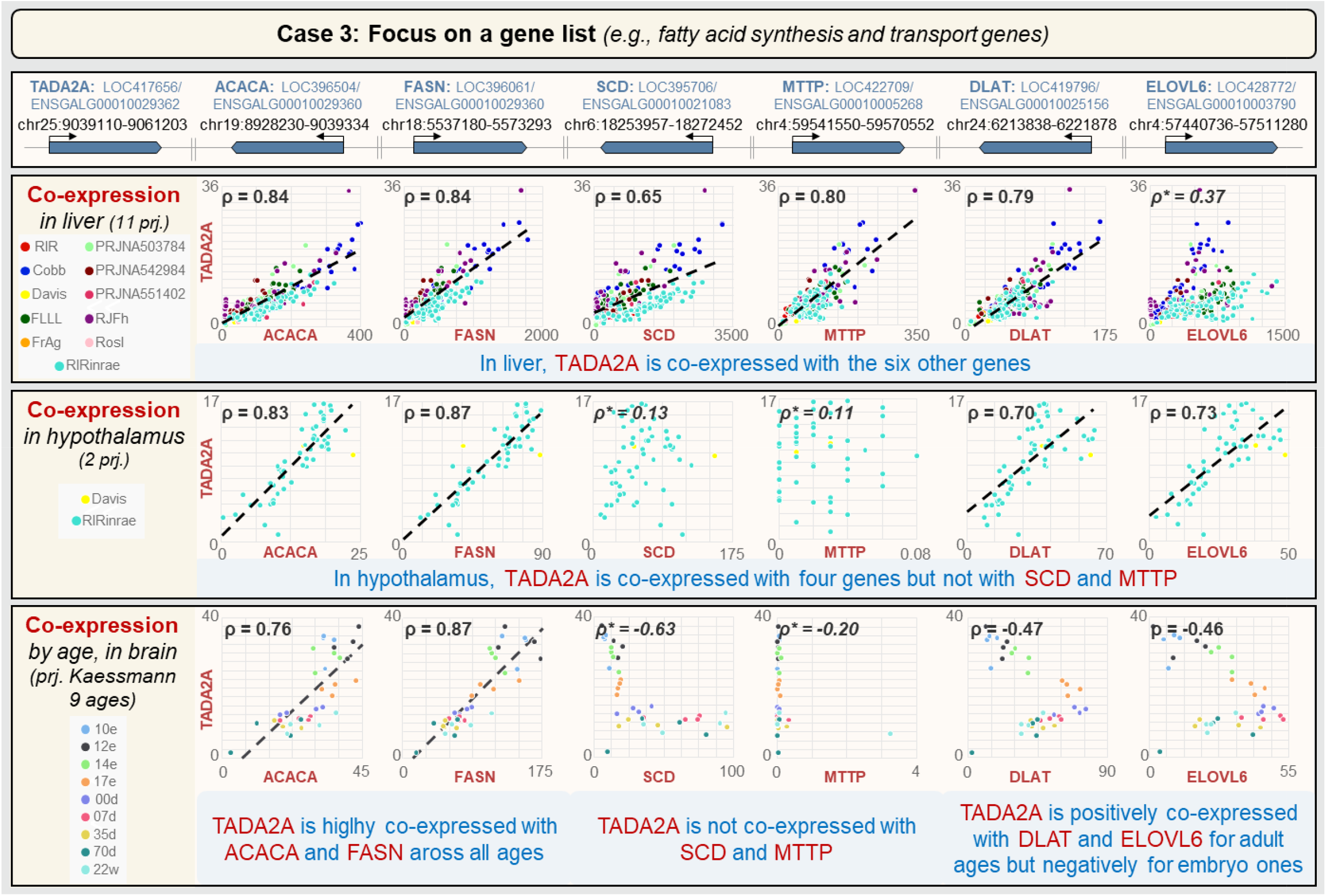
User case 3 – GEGA for exploring a gene list (e.g., fatty acid synthesis and transport genes). The regression line is not shown and the correlation value is indicated in italics and with a (^*^) if the correlation is ≤ |0.2| or if a group effect is observed. Aadditional information on projects is available on GEGA via the hyperlink associated with “projects” in the “Expression Graphical View”. e: embryonic day; d: day; w: week. chr.: chromosome; prj.: project.

## DISCUSSION

The GEGA tool presented in this article integrates and synthesizes the gene models from the reference NCBI-RefSeq and Ensembl/GENCODE databases with those from international collaborative projects such as FAANG and NONCODE (3). Usually, the scientific community interested in chicken genes handle either NCBI-RefSeq or Ensembl/GENCODE identifiers, *i.e*., LOCxx or ENSGALGxx identifiers respectively. One strength of GEGA is that both identifiers can be used in addition to the HGNC name of the gene. However, for gene existing in the two reference databases, the ENSGALGxx gene identifiers used by the user may not correspond to a gene model since the request boxes by default match to the NCBI-RefSeq identifiers. The user has to use the “Function all naming gene” column (that gathers all the gene naming columns) to identify the corresponding gene model. The accumulation of the six annotation databases not only increases the number of gene models but also allows to analyze the repeatability of these models through these databases. Note that it can be difficult to assess the accuracy of the gene models, particularly in terms of exon/intron structure or gene fusions/splits due to different factors as short-sequencing RNA-seq samples, the variety of bioinformatics analysis methods used for the gene modelling or also the impossibility to gather all the condition-specific RNA-seq samples with an enough high sequencing depth. However, the repeated presence of a model at a specific locus in various databases supports the existence of this gene model, even if the transcript isoforms supporting these models are still badly described in farm species (2, 3, 35). Moreover, the observation of an expression in at least one of the 47 tissues is another argument for the existence of a gene model, especially for lncRNAs, which are weakly expressed and sometimes supported by only a few reads. Indeed, a second strength of GEGA is to provide the expression profiles of these genes across 47 tissues using 1,400 samples grouped into 36 datasets representing the diversity of chicken physiological systems, including some bird specific tissues, *e.g*., the different segments of the oviduct that produce the white egg and shell. As a result, 63,513 out of the 78,323 genes (*i.e*., 81%) are observed to be expressed (normalized TPM expression ≥ 0.1) in at least one of the 47 tissues. Note that some neighboring genes in a same strand orientation may correspond to the same gene as already demonstrated in Muret et al. (36). The observation of high co-expression in the 47 tissues and within the various tissues may be an indicator of such a case, thus inviting caution in biological interpretations. This standardized functional annotation greatly facilitates transcriptomic studies in this species by providing a reliable expression atlas reference. This resource is complementary to the chicken EBI-Expression Atlas (37) used by Ensembl which consists of 26 tissues (including 5 embryonic). However, of the 39 projects, all of which refer to previous genome assemblies, only 5 provide easy access to expression data (baseline project), while the remaining 34 projects correspond to expression differentials experiments, in which only FC between conditions are accessible. Note also that 20 projects used Affymetrix technology, which does not include new gene models. Moreover, GEGA’s interactive graphical interface makes it easy to explore this expression datasets according to different tissues combined to physiological conditions as sex and development stages to also analyze tissue specificity, paving the way for new discoveries. The availability of different tissue-specificity indicators, some of which being fairly basic such as fold-change and others more refined such as the tau metrics, allows us to respond to a request from biologists interested in a gene to situate the expression of this gene among all the chicken tissues, considering that the tissue panel in this present study is wide. The possibility to explore co-expression between different genes of interest is also an added value. Although providing a useful basis, gene expressions in GEGA have some limitations such as sometimes fragmentary metadata which limit the possibilities of analysis, *e.g*., for developmental stages or sex that are provided for some tissues only. This limitation may be overcome in the future by adding new projects (which can be suggested through the “Contact” tab): new metadata well defined could easily be added in a coherent and standardized manner using the nfcore “rnaseq” pipeline (38) used with the .*gtf* file associated to this gene-enriched atlas (3). Furthermore, the addition of GO terms and orthologous relationships with mice and humans combined with co-expression between human and chicken species provides valuable functional insights into the interpretation of genomic data. The chicken is a model species for many human diseases, and comparisons of expression profiles between species provide a wealth of information. Similarly, analysis of co-expression networks between coding and non-coding genes highlights potential transcriptional regulations and opens up promising research prospects. However, current reference databases only provide orthologous relationships between PCGs and a few miRNAs, information that have been integrated in GEGA. The integration of lncRNA orthology into GEGA will therefore be carried out in parallel with advances in this field. One of the issues with GEGA is maintaining it up to date as genome assemblies evolves, in particular with the production of the complete genome sequence from telomere to telomere (T2T) (39). The choice was made to update the GEGA annotation only when new versions of genome assemblies are used as reference in the NCBI-Refseq and Ensembl/ENCODE databases, while facilitating the transition from one assembly to another and allowing users to work with older versions. For a given assembly (currently GRCg7b), versions of the reference gene model annotations are fixed (V106 for NCBI-RefSeq, the latest available) and V107 for Ensembl/GENCODE, equivalent to the actual version v110). In conclusion, despite these limitations, GEGA already provides a powerful analytical framework for functional exploration of the chicken genome. Coupled with the latest advances in genome assembly and genome annotation in terms of gene models, this bioinformatics tool supports the functional characterization of the transcriptome of this model species and paves the way for a refined characterization of the functioning of complex genomes.

## DATA AVAILABILITY

The data underlying this article are available through the GEGA website at https://gega.sigenae.org/. The raw data are also available through the FR-AgENCODE website at https://www.fragencode.org/lnchickenatlas.html.

## FUNDING

This project is funded by the European Union’s Horizon 2020 research and innovation program under grant agreement N°101000236 (GEroNIMO) and by ANR CE20 under ‘EFFICACE’ program. Fabien Degalez is a Ph.D. student supported by the Brittany region (France) and the INRAE (Animal Genetics Division).

## ACKNOWLEDGEMENTS

No acknowledgement is provided.

## SUPPLEMENTARY DATA

No supplementary data is provided.

## AUTHOR CONTRIBUTIONS

No author contributions is provided.

## CONFLICT OF INTEREST

The authors have no conflicts of interest to declare that are relevant to the content of this article.

---

**Supplementary Figure 1.**
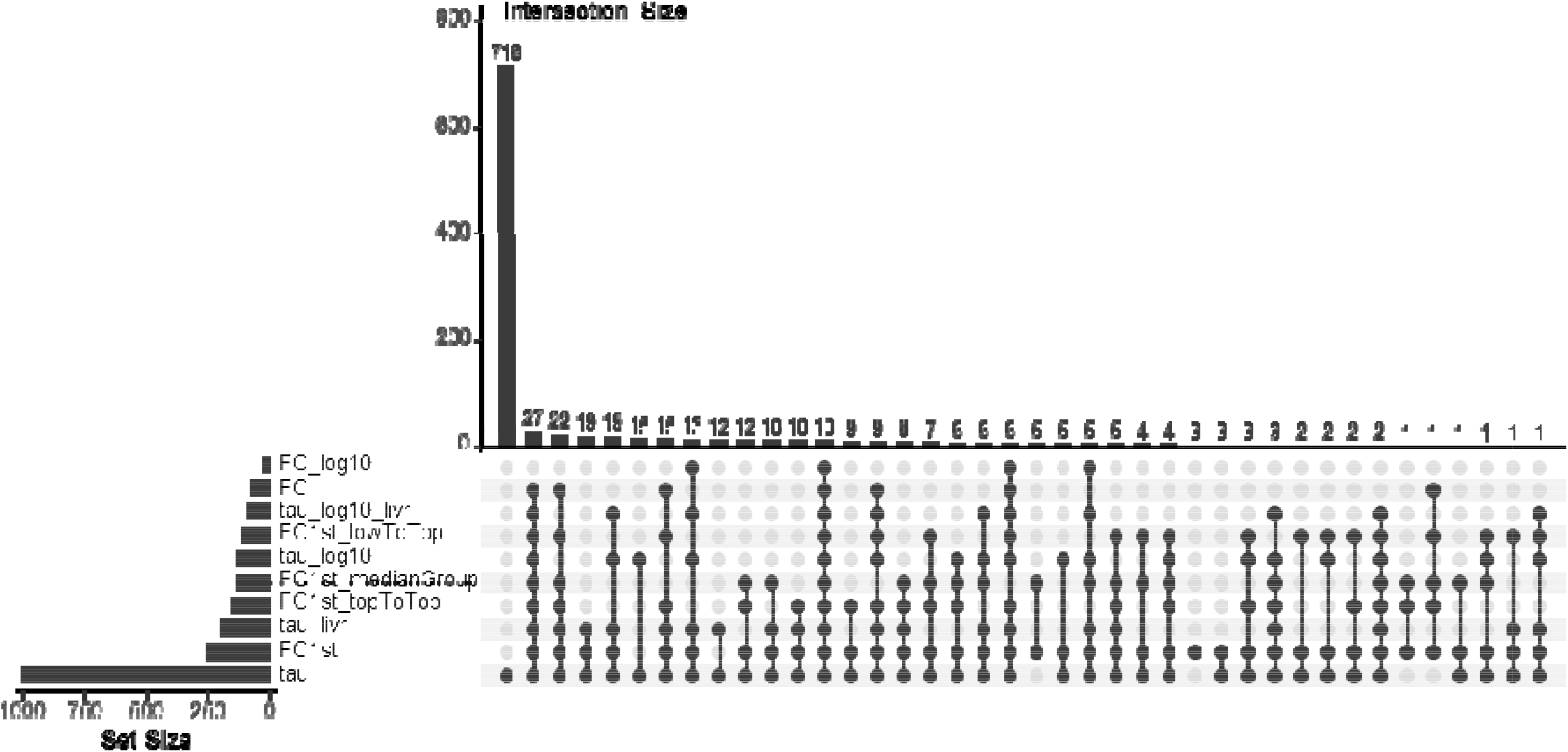
Intersection of the number of tissue-specific PCGs in the liver detected by the different tissue-specific metrics available in GEGA. The notation used for tissue-specific metrics are available in Figure 2.

## Notes

https://gega.sigenae.org/

### Competing Interest Statement

The authors have declared no competing interest.

## REFERENCES

1. Burt, D.W. (2007) Emergence of the chicken as a model organism: implications for agriculture and biology. Poult Sci, 86, 1460–1471.

2. Lagarrigue, S., Lorthiois, M., Degalez, F., Gilot, D. and Derrien, T. (2022) LncRNAs in domesticated animals: from dog to livestock species. Mamm Genome, 33, 248–270.

3. Degalez, F., Charles, M., Foissac, S., Zhou, H., Guan, D., Fang, L., Klopp, C., Allain, C., Lagoutte, L., Lecerf, F., et al. (2023) Enriched atlas of lncRNA and protein-coding genes for the GRCg7b chicken assembly and its functional annotation across 47 tissues. 10.1101/2023.08.18.553750.

4. Morales, J., Pujar, S., Loveland, J.E., Astashyn, A., Bennett, R., Berry, A., Cox, E., Davidson, C., Ermolaeva, O., Farrell, C.M., et al. (2022) A joint NCBI and EMBL-EBI transcript set for clinical genomics and research. Nature, 604, 310–315.

5. Andersson, L., Archibald, A.L., Bottema, C.D., Brauning, R., Burgess, S.C., Burt, D.W., Casas, E., Cheng, H.H., Clarke, L., Couldrey, C., et al. (2015) Coordinated international action to accelerate genome-to-phenome with FAANG, the Functional Annotation of Animal Genomes project. Genome Biology, 16, 57.

6. Giuffra, E., Tuggle, C.K., and FAANG Consortium (2019) Functional Annotation of Animal Genomes (FAANG): Current Achievements and Roadmap. Annu Rev Anim Biosci, 7, 65–88.

7. Tixier-Boichard, M., Fabre, S., Dhorne-Pollet, S., Goubil, A., Acloque, H., Vincent-Naulleau, S., Ross, P., Wang, Y., Chanthavixay, G., Cheng, H., et al. (2021) Tissue Resources for the Functional Annotation of Animal Genomes. Frontiers in Genetics, 12.

8. Zhao, L., Wang, J., Li, Y., Song, T., Wu, Y., Fang, S., Bu, D., Li, H., Sun, L., Pei, D., et al. (2021) NONCODEV6: an updated database dedicated to long non-coding RNA annotation in both animals and plants. Nucleic Acids Res, 49, D165–D171.

9. NCBI-RefSeq (2021) bGalGal1.mat.broiler.GRCg7b - Genome - Assembly - NCBI. NCBI.

10. NCBI-RefSeq (2022) bGalGal1.mat.broiler.GRCg7b - Genome - Annotation - NCBI v106.

11. EMBL EBI Ensembl/GENCODE (2022) bGalGal1.mat.broiler.GRCg7b - Genome - Annotation - Ensembl v107.

12. NCBI-RefSeq (2023) bGalGal1.mat.broiler.GRCg7b - Genome - Annotation Report - NCBI v106.

13. EMBL EBI Ensembl/GENCODE (2023) bGalGal1.mat.broiler.GRCg7b - Genome – Annotation Report Samples - Ensembl v107.

14. Foissac, S., Djebali, S., Munyard, K., Vialaneix, N., Rau, A., Muret, K., Esquerré, D., Zytnicki, M., Derrien, T., Bardou, P., et al. (2019) Multi-species annotation of transcriptome and chromatin structure in domesticated animals. BMC Biology, 17, 108.

15. Kern, C., Wang, Y., Chitwood, J., Korf, I., Delany, M., Cheng, H., Medrano, J.F., Van Eenennaam, A.L., Ernst, C., Ross, P., et al. (2018) Genome-wide identification of tissue-specific long non-coding RNA in three farm animal species. BMC Genomics, 19, 684.

16. Jehl, F., Muret, K., Bernard, M., Boutin, M., Lagoutte, L., Désert, C., Dehais, P., Esquerré, D., Acloque, H., Giuffra, E., et al. (2020) An integrative atlas of chicken long non-coding genes and their annotations across 25 tissues. Sci Rep, 10, 20457.

17. NCBI-RefSeq (2022) Coordinate remapping service: NCBI.

18. Lizio, M., Deviatiiarov, R., Nagai, H., Galan, L., Arner, E., Itoh, M., Lassmann, T., Kasukawa, T., Hasegawa, A., Ros, M.A., et al. (2017) Systematic analysis of transcription start sites in avian development. PLOS Biology, 15, e2002887.

19. Quinlan, A.R. and Hall, I.M. (2010) BEDTools: a flexible suite of utilities for comparing genomic features. Bioinformatics, 26, 841–842.

20. Yanai, I., Benjamin, H., Shmoish, M., Chalifa-Caspi, V., Shklar, M., Ophir, R., Bar-Even, A., Horn-Saban, S., Safran, M., Domany, E., et al. (2005) Genome-wide midrange transcription profiles reveal expression level relationships in human tissue specification. Bioinformatics, 21, 650–659.

21. Wucher, V., Legeai, F., Hédan, B., Rizk, G., Lagoutte, L., Leeb, T., Jagannathan, V., Cadieu, E., David, A., Lohi, H., et al. (2017) FEELnc: a tool for long non-coding RNA annotation and its application to the dog transcriptome. Nucleic Acids Res, 45, e57.

22. Smedley, D., Haider, S., Ballester, B., Holland, R., London, D., Thorisson, G. and Kasprzyk, A. (2009) BioMart – biological queries made easy. BMC Genomics, 10, 22.

23. Ashburner, M., Ball, C.A., Blake, J.A., Botstein, D., Butler, H., Cherry, J.M., Davis, A.P., Dolinski, K., Dwight, S.S., Eppig, J.T., et al. (2000) Gene Ontology: tool for the unification of biology. Nat Genet, 25, 25–29.

24. The Gene Ontology Consortium, Aleksander, S.A., Balhoff, J., Carbon, S., Cherry, J.M., Drabkin, H.J., Ebert, D., Feuermann, M., Gaudet, P., Harris, N.L., et al. (2023) The Gene Ontology knowledgebase in 2023. Genetics, 224, iyad031.

25. Nicholas, F., Tammen, I. and Hub, S.I. (1995) Online Mendelian Inheritance in Animals (OMIA).

26. GTEx Consortium (2023) GTEx Portal-Bulk tissue expression.

27. Cardoso-Moreira, M., Halbert, J., Valloton, D., Velten, B., Chen, C., Shao, Y., Liechti, A., Ascenção, K., Rummel, C., Ovchinnikova, S., et al. (2019) Gene expression across mammalian organ development. Nature, 571, 505–509.

28. Derrien, T., Johnson, R., Bussotti, G., Tanzer, A., Djebali, S., Tilgner, H., Guernec, G., Martin, D., Merkel, A., Knowles, D.G., et al. (2012) The GENCODE v7 catalog of human long noncoding RNAs: Analysis of their gene structure, evolution, and expression. Genome Res., 22, 1775–1789.

29. Chapman, D.L., Garvey, N., Hancock, S., Alexiou, M., Agulnik, S.I., Gibson-Brown, J.J., Cebra-Thomas, J., Bollag, R.J., Silver, L.M. and Papaioannou, V.E. (1996) Expression of the T-box family genes, Tbx1-Tbx5, during early mouse development. Dev Dyn, 206, 379–390.

30. Hatcher, C., Kim, M., Maha, C., Goldstein, M., Wong, B., Mikawa, T. and Basson, C. (2001) TBX5 transcription factor regulates cell proliferation during cardiogenesis. Developmental biology, 230, 177–188.

31. Hori, Y., Tanimoto, Y., Takahashi, S., Furukawa, T., Koshiba-Takeuchi, K. and Takeuchi, J.K. (2018) Important cardiac transcription factor genes are accompanied by bidirectional long non-coding RNAs. BMC Genomics, 19, 967.

32. Douaud, M., Feve, K., Pituello, F., Gourichon, D., Boitard, S., Leguern, E., Coquerelle, G., Vieaud, A., Batini, C., Naquet, R., et al. (2011) Epilepsy Caused by an Abnormal Alternative Splicing with Dosage Effect of the SV2A Gene in a Chicken Model. PLoS One, 6, e26932.

33. Desert, C., Baéza, E., Aite, M., Boutin, M., Le Cam, A., Montfort, J., Houee-Bigot, M., Blum, Y., Roux, P.F., Hennequet-Antier, C., et al. (2018) Multi-tissue transcriptomic study reveals the main role of liver in the chicken adaptive response to a switch in dietary energy source through the transcriptional regulation of lipogenesis. BMC Genomics, 19, 187.

34. Hamilton, J.A., Hillard, C.J., Spector, A.A. and Watkins, P.A. (2007) Brain uptake and utilization of fatty acids, lipids and lipoproteins: application to neurological disorders. J Mol Neurosci, 33, 2–11.

35. Smith, J., Alfieri, J.M., Anthony, N., Arensburger, P., Athrey, G.N., Balacco, J., Balic, A., Bardou, P., Barela, P., Bigot, Y., et al. (2022) Fourth Report on Chicken Genes and Chromosomes 2022. Cytogenet Genome Res, 162, 405–528.

36. Muret, K., Klopp, C., Wucher, V., Esquerré, D., Legeai, F., Lecerf, F., Désert, C., Boutin, M., Jehl, F., Acloque, H., et al. (2017) Long noncoding RNA repertoire in chicken liver and adipose tissue. Genetics, Selection, Evolution⍰: GSE, 49.

37. Moreno, P., Fexova, S., George, N., Manning, J.R., Miao, Z., Mohammed, S., Muñoz-Pomer, A., Fullgrabe, A., Bi, Y., Bush, N., et al. (2022) Expression Atlas update: gene and protein expression in multiple species. Nucleic Acids Research, 50, D129–D140.

38. Patel, H., Ewels, P., Peltzer, A., Botvinnik, O., Manning, J., Sturm, G., Garcia, M.U., Moreno, D., Vemuri, P., bot, nf-core, et al. (2023) nf-core/rnaseq: nf-core/rnaseq v3.13.2 - Cobalt Colt. 10.5281/zenodo.10171269.

39. Huang, Z., Xu, Z., Bai, H., Huang, Y., Kang, N., Ding, X., Liu, J., Luo, H., Yang, C., Chen, W., et al. (2023) Evolutionary analysis of a complete chicken genome. Proceedings of the National Academy of Sciences, 120, e2216641120.

